# The Cell Division Protein FzlA Performs a Conserved Function in Diverse Alphaproteobacteria

**DOI:** 10.1101/2024.05.29.596507

**Authors:** Isaac P. Payne, Brody Aubry, Jordan M. Barrows, Pamela J.B. Brown, Erin D. Goley

**Affiliations:** Department of Biological Chemistry, Johns Hopkins University School of Medicine, Baltimore, Maryland, USA; Division of Biological Sciences, University of Missouri, Columbia, Missouri, USA; Department of Integrative Structural and Computational Biology, Scripps Research Institute, La Jolla, California, USA

## Abstract

In almost all bacteria, the tubulin-like GTPase FtsZ polymerizes to form a “Z-ring” that marks the site of division. FtsZ recruits other proteins, collectively known as the divisome, that together remodel and constrict the envelope. Constriction is driven by peptidoglycan (PG) cell wall synthesis by the glycosyltransferase FtsW and the transpeptidase FtsI (FtsWI), but these enzymes require activation to function. How recruitment of FtsZ to the division site leads to FtsWI activation and constriction remains largely unknown. Previous work in our laboratory demonstrated that an FtsZ-binding protein, FzlA, is essential for activation of FtsWI in the alphaproteobacterium *Caulobacter crescentus*. Additionally, we found that FzlA also binds to a DNA translocase called FtsK, suggesting that it may link constriction activation to chromosome segregation. FzlA is conserved throughout alphaproteobacteria but has only been examined in detail in *C. crescentus*. Here, we explored whether FzlA function is conserved in diverse alphaproteobacteria. We assessed FzlA homologs from *Rickettsia parkeri* and *Agrobacterium tumefaciens*, and found that, similar to *C. crescentus* FzlA, they bind directly to FtsZ and localize to midcell. The FtsZ-FzlA interaction interface is conserved, as we demonstrated that FzlA from each of the three species examined can bind to FtsZ from any of the three *in vitro*. Additionally, we determined that *A. tumefaciens* FzlA can fulfill the essential function of FzlA when produced in *C. crescentus*, indicating conservation of function. These results suggest that FzlA serves as an important regulator that coordinates chromosome segregation with envelope constriction across diverse alphaproteobacteria.

**Importance:** Cell division is essential for bacterial replication and must be highly regulated to ensure robust remodeling of the cell wall in coordination with segregation of the genome to daughter cells. In *Caulobacter crescentus*, FzlA plays a major role in regulating this process by activating cell wall synthesis in a manner that couples constriction to chromosome segregation. FzlA is broadly conserved in alphaproteobacteria suggesting it plays a similar function across this class of bacteria. Here we have shown that, indeed, FzlA biochemical interactions and function are conserved in diverse alphaproteobacteria. Because FzlA is conserved in alphaproteobacterial human pathogens, understanding this protein and its interactome could present therapeutic benefits by identifying potential antibiotic targets to treat infections.

## Introduction

Bacterial cell division is a tightly regulated process that allows bacteria to propagate in nearly every corner of the world. FtsZ, a dynamic tubulin-like GTPase, regulates cell division by polymerizing into a ring (the Z-ring) at the site of division and recruiting other division-associated proteins (collectively known as the divisome) (1). Together, divisome proteins remodel and constrict the cell envelope in coordination with other cell cycle events like chromosome replication and segregation.

The cell envelope in Gram-negative bacteria consists of a cell wall placed between an inner and outer membrane. The cell wall is made of peptidoglycan (PG), a meshwork of glycan strands crosslinked by short peptides, and is responsible for maintaining cell shape and providing structural integrity (2). For cells to divide, they must remodel their cell envelope through synthesis of new cell wall, which is performed by the glycosyltransferase FtsW and transpeptidase FtsI during division. FtsW and FtsI work in a complex (FtsWI) and must be activated to drive constriction (1, 3–5). Bacterial division requires precise spatiotemporal control of constriction to ensure that each daughter cell receives an undamaged copy of the genome while maintaining cell envelope integrity throughout constriction and cell separation. However, our understanding of FtsWI activation is incomplete.

To study regulation of cell division, we use *Caulobacter crescentus*, a Gram-negative alphaproteobacterium whose cell cycle and morphogenesis mechanisms are well characterized (6). Previously, we identified an FtsZ-binding partner, FzlA, that is conserved throughout alphaproteobacteria and is essential for division in *C. crescentus* and in *Agrobacterium tumefaciens* (7–9). In *C. crescentus*, FzlA-depleted cells are unable to constrict, even though FtsZ assembles into rings in its absence, suggesting a role for FzlA in regulating activity of the divisome (7). Indeed, *fzlA* becomes nonessential when FtsWI are hyperactivated by mutation (8). Moreover, single molecule tracking studies demonstrated that FzlA signals to convert FtsWI to a slow-moving, active state, and that FzlA molecules can move in a complex with active FtsWI (10). Together, these data indicate that FzlA lies in a signaling pathway leading to activation of FtsWI for constriction.

FzlA forms a homodimer, and mutagenesis studies revealed that it possesses at least two essential surfaces (9). One surface is required for binding FtsZ: when residues in this region are mutated, FzlA fails to interact with FtsZ *in vitro* and cannot localize or support division in cells (9). The second essential surface comprises the C-terminal tail of FzlA, which is highly conserved across FzlA homologs. The C-terminal tail is required to bind FtsK, a cell division protein and DNA translocase that segregates the chromosomal terminus and promotes dimer resolution at the end of DNA replication (10). Disruption of FzlA-FtsK-mediated regulation of FtsWI leads to DNA damage and cell death (10). We therefore propose that FzlA interacts with FtsK to sense clearance of the chromosome from the division plane and regulate activation of FtsWI to ensure constriction does not occur prior to completion of chromosome segregation.

FzlA is conserved throughout alphaproteobacteria, a group described as the “Darwin’s finches of bacteria” (11) because its members display a wide range of lifestyles, morphologies, and genome sizes. While FzlA has been characterized in *C. crescentus*, we sought to understand if this conserved protein performs a similar function in other alphaproteobacteria with distinct lifestyles, including in plant and human pathogens. To this end, we selected the alphaproteobacteria *Agrobacterium tumefaciens* and *Rickettsia parkeri* to explore whether FzlA activities are conserved. Both of these bacteria encode FzlA homologs, but they have very different lifestyles and growth mechanisms. *A. tumefaciens* is a plant pathogen in the order Hyphomicrobiales that undergoes polar growth, but still divides at midcell (12, 13). *R. parkeri* is a tick-borne, obligate intracellular human pathogen in the order Rickettsialles that has a streamlined genome(14, 15) and lacks a number of other divisome proteins conserved in *C. crescentus*. We reasoned that examining FzlA biochemistry and function from these two additional alphaproteobacteria should reveal if FzlA has a broadly conserved function.

Here we analyze the function of FzlA from *A. tumefaciens* and *R. parkeri* and compare them to that of *C. crescentus* FzlA. We find that the FtsZ-FzlA interaction interface is conserved based on structural predictions and biochemical evidence. We also observe that FzlA from *A. tumefaciens* and *R. parkeri* can localize to midcell in *C. crescentus* and that cells producing *A. tumefaciens* FzlA instead of *C. crescentus* FzlA are viable, suggesting at least partial conservation of function. These results demonstrate that FzlA plays an important role regulating cell division across alphaproteobacteria, including bacteria that pose threats to human health.

## Results

### AlphaFold predicts a conserved interface between FtsZ and FzlA

The ability of FzlA to interact with FtsZ is essential for division in *C. crescentus* (7). While this interaction has been well characterized biochemically, there is currently no structural detail of a FtsZ-FzlA complex (7–9). Before characterizing the function of FzlA in other alphaproteobacteria, we sought to generate a structural model of *C. crescentus* FzlA bound to FtsZ. To do this, we used ColabFold (16) to generate a model of an interaction between monomers of *C. crescentus* FzlA (CcFzlA) and FtsZ (CcFtsZ) (Fig. 1A). The predicted aligned error (PAE) for intramolecular alignment of residues within FtsZ or FzlA is low, with the exception of the disordered linker region of FtsZ (Supp. Fig. 1A). Importantly, the PAEs for residues in the interface between FzlA and FtsZ are low, suggesting a high confidence model (Supp. Fig. 1A). We mapped CcFzlA residues known to be involved in the interaction with CcFtsZ from our previous work onto our model (Fig. 1A, residues in red). All of these residues on CcFzlA are located at or near the interface between CcFzlA and CcFtsZ in our predicted model, supporting the validity of the model. We identified several residues on CcFtsZ (Q170, E178, N250) that were predicted to interact with CcFzlA, with some of these residues having a predicted interaction with residues on CcFzlA previously implicated in interaction with FtsZ (9). Almost all of the CcFtsZ residues identified as interacting with CcFzlA are conserved in alphaproteobacterial FtsZs, while they are divergent in FtsZs from other groups of bacteria (Table 1). This suggests that this model accurately identified residues involved in the alphaproteobacterial-specific FtsZ-FzlA interaction.

**Figure 1.**
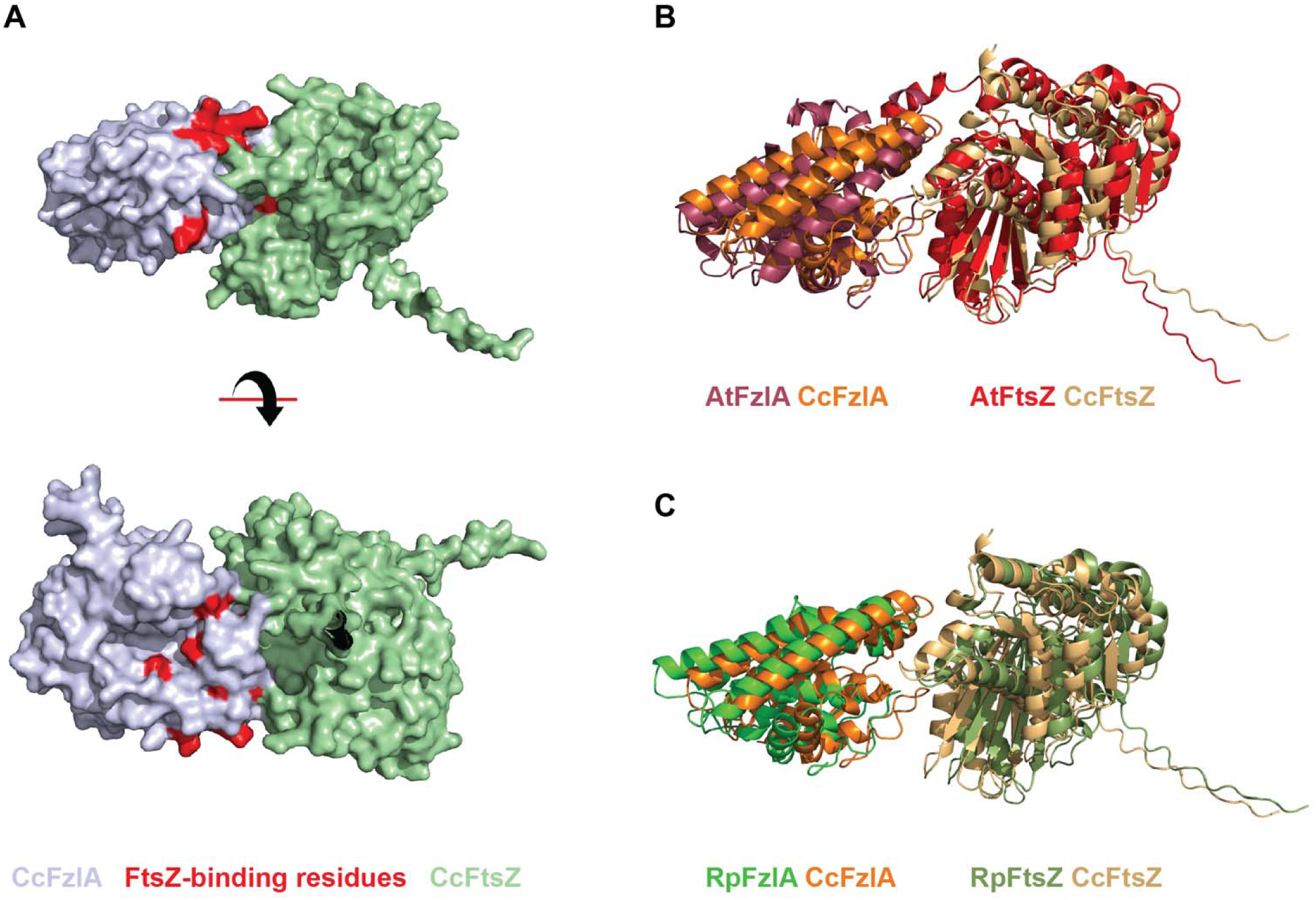
The FtsZ-FzlA binding interface is conserved in alphaproteobacteria. **A.** Surface plot of the predicted interaction between FzlA (gray) and FtsZ (green) generated using ColabFold. FzlA residues that have been previously shown to be essential for the FtsZ interaction are shown in red. **B-C.** The predicted ColabFold structure of either the *A. tumefaciens* FtsZ-FzlA interaction or the *R. parkeri* FtsZ-FzlA interaction, each overlaid with the predicted structure for *C. crescentus* FtsZ and FzlA.

**Table 1.** Conservation of FtsZ residues predicted to be involved in the FtsZ-FzlA interaction in *C. crescentus*. Alphaproteobacterial species are shown in blue while bacteria from other classes of bacteria are shown in orange. The CcFtsZ residues were identified using the model of FtsZ-FzlA generated using ColabFold. Identical residues to the CcFtsZ residue are shown in green. Conserved, but nonidentical residues are shown in yellow. Residues that are not conserved are shown in red.

| CcFtsZ Residue | *R. parkeri* | *A. tumefaciens* | *M. magneticum* | *S. paucimobilis* | *R. capsulatus* | *E. coli* | *M. tuberculosis* | *B. subtilis* | *B. burgdorferi* | *N. meningitidis* |
| --- | --- | --- | --- | --- | --- | --- | --- | --- | --- | --- |
| Q170 | Q | Q | Q | Q | Q | D | D | D | Q | D |
| R174 | R | R | R | L | R | K | Q | E | T | T |
| A176 | A | A | A | A | A | L | G | V | V | L |
| E178 | E | D | E | A | E | R | A | K | K | E |
| N246 | S | A | A | K | K | M | I | K | S | Q |
| N250 | N | N | N | N | N | S | S | S | N | S |
| E255 | H | E | D | G | E | D | - | - | E | D |

As a first step in determining if this binding interface is conserved in other alphaproteobacteria, we generated models using ColabFold of the FtsZ-FzlA interaction in *A. tumefaciens* and *R. parkeri*. We aligned the top-ranked FtsZ-FzlA models from both *A. tumefaciens* and *R. parkeri* with our *C. crescentus* model, and noted that the overall orientation of FzlA and FtsZ in each species was the same (Fig. 1A,B). However, unlike for the CcFzlA-CcFtsZ model, the PAEs for residues in the FzlA-FtsZ interface in *A. tumefaciens* and *R. parkeri* predicted complexes indicated low confidence (Supp. Fig. 1B,C). Perhaps accordingly, we determined RMSD values of 4.905 and 4.627 Å for *C. crescentus* vs *A. tumefaciens* and *R. parkeri* complexes, respectively. The low confidence of the *A. tumefaciens* and *R. parkeri* models required us to further explore potential conservation of this interaction biochemically and genetically.

### FtsZ polymerization properties are similar across alphaproteobacteria

Our structural predictions motivated us to directly test if the FtsZ-FzlA interaction is conserved in alphaproteobacteria using purified proteins *in vitro*. Our binding assays rely on the ability of FtsZ to polymerize, so we first sought to confirm that FtsZs from the selected alphaproteobacteria have similar polymerization properties. While the polymerization properties of FtsZ from *C. crescentus* (CcFtsZ) and *A. tumefaciens* (AtFtsZ2) are known (17, 18), *R. parkeri* FtsZ (RpFtsZ) has never been characterized. Using negative stain transmission electron microscopy (TEM), we analyzed filaments formed by the three FtsZs incubated in the presence of GTP to induce polymerization. Each of the FtsZs tested readily polymerized under these conditions, forming mostly single or paired protofilaments that are gently curved (Fig. 2A).

**Figure 2.**
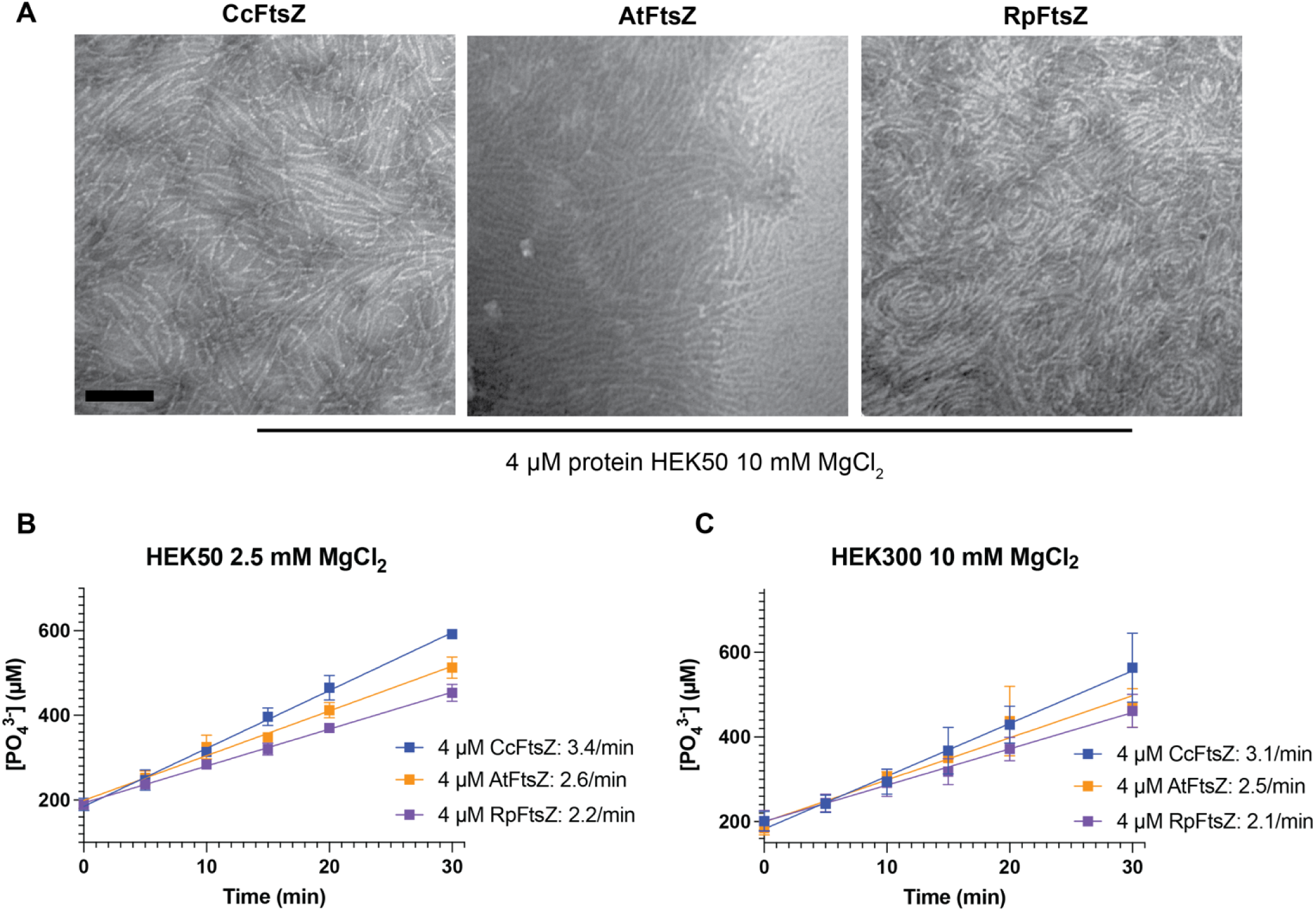
Alphaproteobacterial FtsZs form similar polymers and have similar GTPase rates. **A.** Negative stain TEM images of 4 µM FtsZ under polymerizing conditions from each of the alphaproteobacteria tested. Scale bar = 100 nm. **B-C.** Plots of inorganic phosphate concentration over time for the indicated proteins and concentrations. Apparent GTPase rates per molecule of FtsZ for each condition are indicated. Mean and SEM for three replicates are shown. HEK – Hepes, EDTA, KCl; number refers to KCl concentration in mM.

To understand the dynamic properties of alphaproteobacterial FtsZs, we next measured the GTPase rates of CcFtsZ, AtFtsZ2, and RpFtsZ using a malachite green assay to monitor the release of inorganic phosphate (P_i_) over time. For CcFtsZ and AtFtsZ2 we observed GTPase rates of about 3 and 2.5 P_i_ released per minute per FtsZ molecule, respectively, at both low and high salt concentrations (Fig. 2B,C). These rates are consistent with previously measured rates for CcFtsZ and AtFtsZ2 (17, 18). For RpFtsZ, we measured a GTPase rate of 2.1 P_i_ released per minute per FtsZ molecule (Fig. 2B,C). We conclude that alphaproteobacterial FtsZs exhibit similar polymerization properties *in vitro*.

### The FtsZ-FzlA interaction is conserved in *A. tumefaciens* and *R. parkeri*

With purified FtsZs in hand, we next sought to directly test interactions with FzlA for each species. To do this, we used a previously established high speed co-pelleting assay in which FtsZ polymers pellet and can bring bound FzlA into the pellet to assess binding (7). If FtsZ polymers are not present, FzlA is found in the supernatant. Using purified FtsZ and FzlA from *C. crescentus*, *R. parkeri*, and *A. tumefaciens*, we assessed the ability of each FtsZ to bind to its cognate FzlA. In agreement with previous studies, we saw a significant increase in the amount of CcFzlA in the pellet in the presence of polymerized CcFtsZ (Fig. 3A,B). For RpFzlA and RpFtsZ we observed a similar increase in the amount of FzlA in the pellet in the presence of FtsZ polymers, suggesting that the FtsZ-FzlA interaction is maintained in *R. parkeri* (Fig. 3A,B). Interestingly, *A. tumefaciens* encodes multiple FtsZs, with AtFtsZ2 being most similar to *C. crescentus* and *R. parkeri* FtsZ (17). AtFzlA interacted with AtFtsZ2, as we saw an increase in the amount of FzlA in the pellet under polymerizing conditions (Fig. 3A,B). We noticed that the increase of AtFzlA in the pellet was not as large as that for *C. crescentus* and *R. parkeri* and reasoned that this is likely because AtFtsZ2 polymers do not pellet well on their own (Supp. Fig. 2A,B), limiting the amount of AtFzlA that can co-pellet. These results suggest that the FtsZ-FzlA interaction is conserved in *A. tumefaciens* and *R. parkeri*.

**Figure 3.**
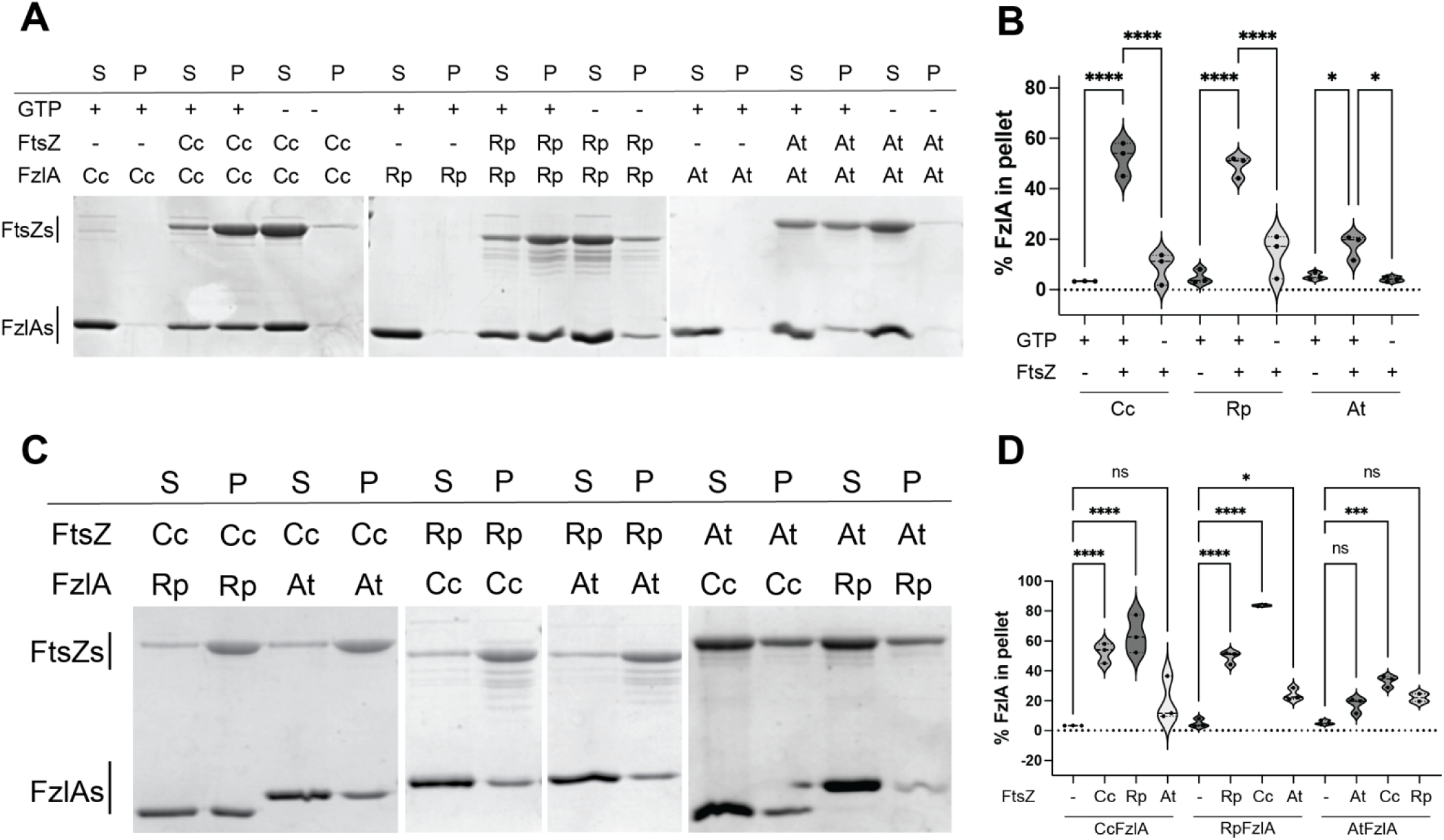
The FtsZ-FzlA interaction is conserved *in vitro*. **A-B**. Co-pelleting of purified FzlA and its cognate FtsZ from three different alphaproteobacteria. Representative images of the Coomassie blue SDS-PAGE gels from each condition are shown. Using the Coomassie-blue SDS-PAGE gels, the amount of FzlA in the pellet for each condition was quantified and plotted. Each condition was performed in triplicate. **C-D.** Co-pelleting of purified FzlA and its non-cognate FtsZ from three different alphaproteobacteria. Representative images of the Coomassie blue SDS-PAGE gels from each condition are shown. The amount of FzlA in the pellet for each condition was quantified and plotted. Each experiment was performed in triplicate. For each quantification, a one-way ANOVA with Šídák’s multiple comparison test was performed to assess differences between the conditions tested. *P < 0.0332, ***P < 0.0002, ****P < 0.0001.

In addition to AtFtsZ2, *A. tumefaciens* encodes two additional FtsZ homologs, AtFtsZ1 and AtFtsZ3 (19). AtFtsZ3 is significantly truncated, bearing only half of the GTPase domain, while AtFtsZ1 contains the full GTPase domain, but lacks the C-terminal linker and conserved C-terminal peptide found in canonical FtsZ homologs (19). We previously showed that AtFtsZ1 cannot polymerize on its own but can co-polymerize with AtFtsZ2 (17). We therefore assessed whether AtFzlA could associate with AtFtsZ1 or with AtFtsZ1-AtFtsZ2 co-polymers. However, we did not observe AtFzlA binding with AtFtsZ1 polymers. Although AtFzlA co-pelleted with AtFtsZ1-AtFtsZ2, there was no difference in the amount of AtFzlA that pelleted with co-polymers compared to AtFtsZ2 alone, suggesting that AtFtsZ2 is sufficient for FzlA interaction (Supp. Fig. 2C,D).

The above results indicated that FzlA and FtsZ directly interact in diverse alphaproteobacteria. We next wanted to determine whether FzlA could bind FtsZ from a different bacterial species, which would suggest that specific residues involved in this interaction are conserved. To do this, we performed high speed co-pelleting assays, as described previously, with different combinations of FtsZ and FzlA from the three alphaproteobacterial species. We observed that CcFzlA binds to RpFtsZ and AtFtsZ2, as we saw an increase of CcFzlA in the pellet compared to the condition with no FtsZ (Fig. 3C,D). Similarly, when we performed these assays with RpFzlA or AtFzlA, we saw an increase in the amount of FzlA in the pellet when using FtsZ from any of the alphaproteobacterial species (Fig. 3C,D). These data indicate that FzlA can interact with FtsZ from other alphaproteobacteria and suggest that the residues involved in this interaction are conserved across species.

### The roles of invariant residues of FzlA are conserved

After demonstrating conservation of the FzlA-FtsZ interaction across species, we next wanted to directly test whether residues required for the FzlA-FtsZ interaction in *C. crescentus* are required across alphaproteobacteria. A previous study from our lab identified several residues on CcFzlA that were essential for its function (9). Further analysis showed that several of these residues are involved in the FzlA-FtsZ interaction (9). Other essential residues are located on the C-terminus of FzlA and are dispensable for FtsZ binding but necessary for interaction with FtsK (9, 10). We sought to test if mutating analogous residues in AtFzlA or RpFzlA had similar effects on the FtsZ interaction.

We selected two mutant variants of CcFzlA that failed to bind CcFtsZ in high-speed co-pelleting assays, NB1 and NB2, upon which to model corresponding mutant variants in AtFzlA and RpFzlA. In addition, we included a mutant variant in the C-terminus of CcFzlA that retains the ability to bind CcFtsZ (9). When comparing FzlA sequences, we found that only one of the two residues mutated for the NB1 mutant (W37 in *C. crescentus)* is conserved in both *R. parkeri* and *A. tumefaciens* FzlA. The single residue mutated for the NB2 mutant (E119 in *C. crescentus*) is also conserved. We therefore made W35A mutations in RpFzlA and AtFzlA, analogous to the NB1 mutant, and E113K or E117K mutations in RpFzlA and AtFzlA, respectively, analogous to the NB2 mutation. We attempted to purify these variants to perform high speed co-pelleting assays. We found that the E117K mutant protein of AtFzlA was insoluble, precluding biochemical analysis, so we proceeded with the remaining mutant proteins.

We first confirmed our prior observations that the NB1 or NB2 mutant variants of CcFzlA failed to interact with CcFtsZ, as demonstrated by a decrease in the amount of FzlA found in the pellet (Fig. 4A,B). Mutating the essential C-terminal residue (D227K) of CcFzlA did not decrease the fraction of FzlA in the pellet, suggesting it is not required for the FtsZ-FzlA interaction, as expected (Fig. 4A,B) (9). In *R. parkeri* we observed the same trend, with less RpFzlA in the pellet compared to WT for both the W35A and E113K mutants (Fig. 4C,D). As with CcFzlA, mutating the C-terminal tail of RpFzlA (D221K) had no effect on the RpFtsZ-RpFzlA interaction (Fig. 4C,D).

**Figure 4.**
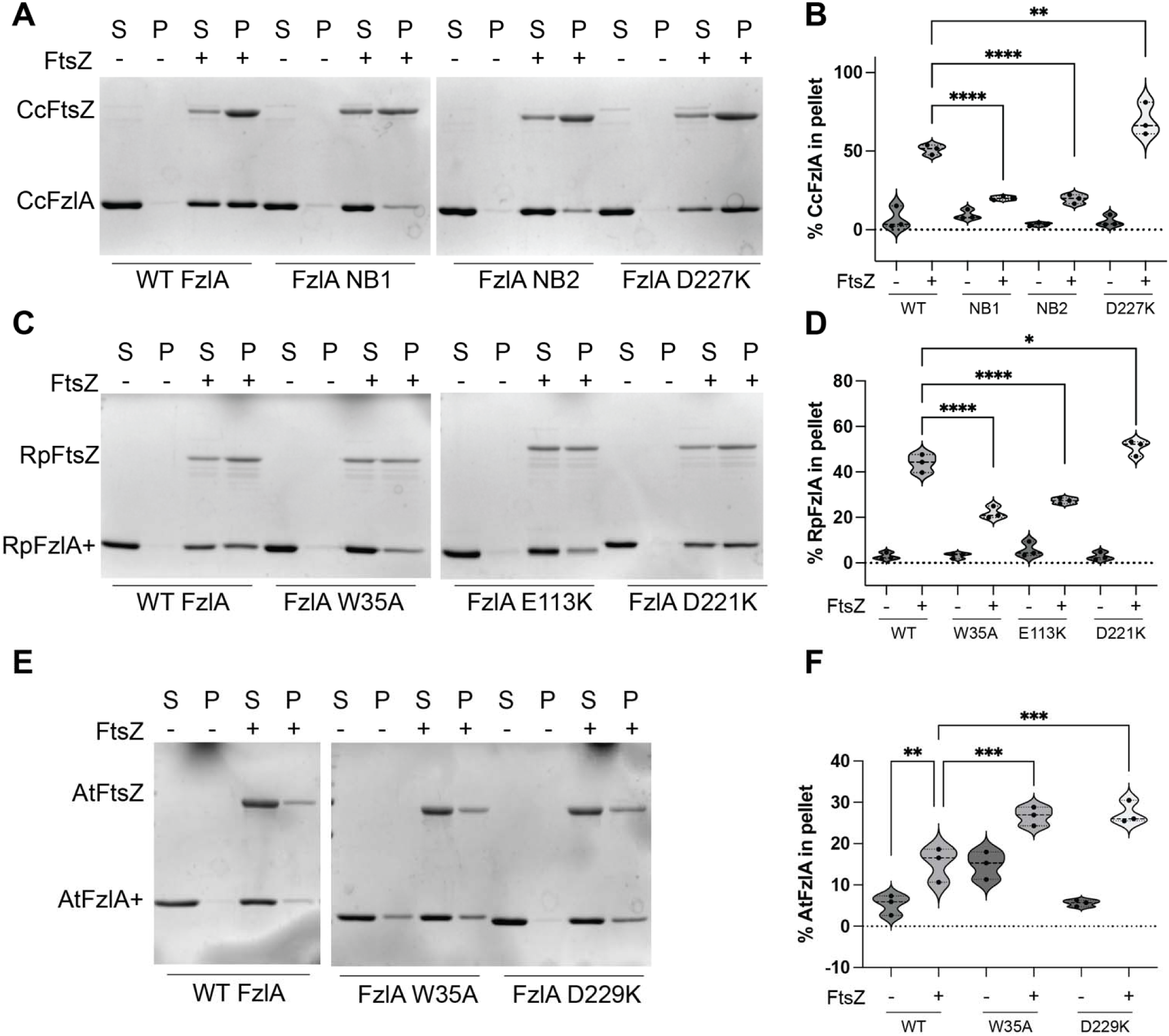
Conserved residues on FzlA participate in FtsZ binding across species. Co-pelleting of purified mutant FzlAs with FtsZ from three alphaproteobacteria. Representative images of the Coomassie blue SDS-PAGE gels from each condition are shown. The amount of FzlA in the pellet for each condition was quantified and plotted. Each condition was performed in triplicate. **A-B.** Co-pelleting of purified mutant CcFzlAs with CcFtsZ. NB1 corresponds to W38A R124D mutations and NB2 corresponds to the E119K mutation. **C-D.** Co-pelleting of purified mutant RpFzlAs with RpFtsZ. **E-F.** Co-pelleting of purified mutant AtFzlAs with AtFtsZ. For each quantification, a one-way ANOVA with Šídák’s multiple comparison test was performed to assess differences between the different conditions tested. *P < 0.0332, **P < 0.0021, ***P < 0.0002, ****P < 0.0001.

The W35A mutation in AtFzlA, did not lead to a decrease in the amount of FzlA in the pellet, suggesting that this mutation did not disrupt the interaction. However, there was a higher percentage of the AtFzlA-W35A in the pellet without FtsZ present compared to WT AtFzlA, indicating that this mutation also affected AtFzlA solubility and complicating interpretation of co-pelleting results (Fig. 4E,F). Consistent with CcFzlA and RpFzlA, we found that mutating the essential C-terminal residue (D229K) of AtFzlA does not disrupt FtsZ binding (Fig. 4E,F). From these experiments we can conclude that specific residues involved in the FtsZ-FzlA interaction are conserved in *C. crescentus* and *R. parkeri* FzlA. Although our analysis of AtFzlA mutants was limited by solubility issues, our finding that the C-terminal tail variant of AtFzlA co-pellets with AtFtsZ is at least consistent with conservation of these interaction interfaces. Taken altogether our co-pelleting data demonstrates that the FtsZ-FzlA interaction is conserved in diverse alphaproteobacteria.

### Activities of some invariant FzlA residues are conserved *in vivo* in *A. tumefaciens*

We next sought to use genetic approaches to determine whether residues important for CcFzlA function *in vivo* are similarly important in other alphaproteobacteria. A lack of appropriate genetic tools prevented us from pursuing this question in *R. parkeri*. We therefore tested if the three AtFzlA mutants described in the previous section were able to support viability and cell division of *A. tumefaciens* (Fig. 5). We did this by introducing the FzlA variants on a plasmid, or an empty vector (EV) control, into a FzlA depletion strain (8). This allowed us to express WT *AtfzlA* from the chromosome in the presence of IPTG or express the *AtfzlA* mutants from the plasmid in the presence of cumate (Fig. 5A,B). We found that all strains grew equally well on solid media in the presence of IPTG. Moreover, when induced with cumate, production of WT AtFzlA supported robust colony formation, while cells bearing the EV control did not grow, as expected (Fig. 5A). We visualized cell morphology of each strain when grown in the absence of cumate and imaged by phase contrast microscopy. As we previously demonstrated (8), growth in the presence of WT AtFzlA yielded cells of normal, rod-shaped morphology while depletion of FzlA (EV control) led to cell elongation and formation of bulges and branches from midcell, indicating division failure (Fig. 5B).

**Figure 5.**
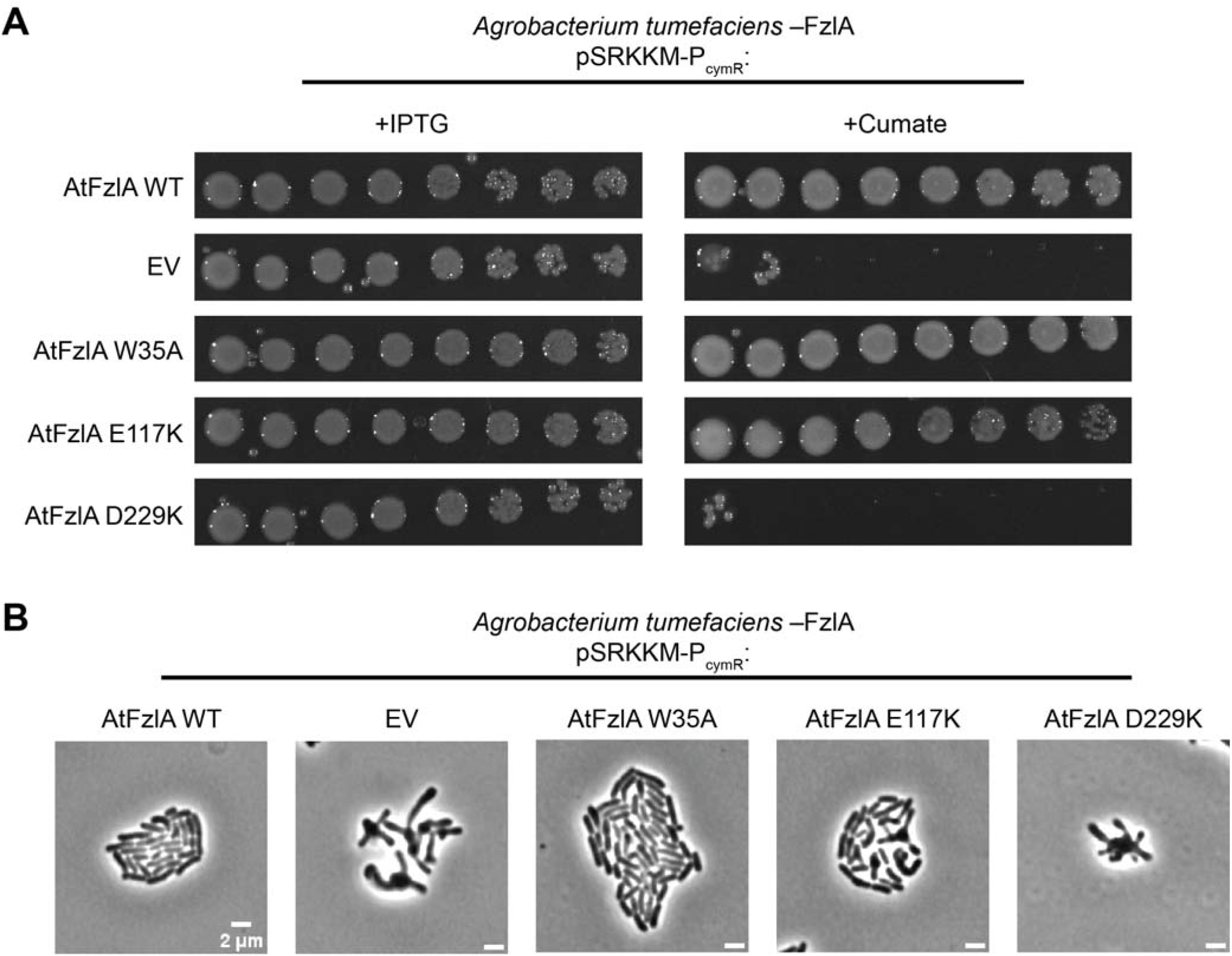
Conserved residues of FzlA are functionally important in *A. tumefaciens*. **A.** Spotting assays of *A. tumefaciens* in a FzlA depletion background with pSRKKM-P*_cymR_* containing *AtfzlA*, empty vector, *AtfzlA-W35A*, *AtfzlA-E117K*, and *AtfzlA-D229K*. When grown on LB Kan with 1mM IPTG, FzlA is expressed from the chromosome. When grown on LB Kan with 0.1 mM cumate, the protein indicated to the left is expressed from the plasmid. **B.** Phase contrast images of cells depleted of AtFzlA for 27 h in liquid ATGN Kan prior to spotting onto ATGN Kan agarose pads with 0.1 mM cumate and incubating for 16 h. Scale bar = 2µm.

Consistent with our findings in *C. crescentus*, production of AtFzlA-E117K resulted in a slight viability defect as evidenced by a decrease in colony forming units (CFUs) compared to cells producing WT AtFzlA (Fig. 5A). There was also an apparent cell division defect when the E117K variant was produced, as assessed by microscopy (Fig. 5B). We observed that the D229K mutant is lethal in *A. tumefaciens* as demonstrated by failure to grow on solid media and profound division failure evident from phase contrast imaging (Fig. 5A,B). This is consistent with our observations in *C. crescentus* that this residue is essential for FzlA function (9, 10).

Surprisingly, we did not observe an effect on viability or morphology when AtFzlA-W35A was produced in *A. tumefaciens.* This mutation was modeled after the NB1 double mutant in CcFzlA W38A R124D (9). The R124 residues is not conserved in *A. tumefaciens fzlA*, so it is possible that mutating the single W35 residue is not sufficient to disrupt the FtsZ-FzlA interaction at this binding interface. This is consistent with our *in vitro* observation that the W35A AtFzlA mutant co-pelleted to some extent with AtFtsZ. Collectively, our biochemical and genetic data suggest that residues we identified as important for CcFzlA function also contribute to FzlA activity and/or function in *A. tumefaciens* and *R. parkeri*.

### A. tumefaciens and R. parkeri FzlA can localize to midcell in C. crescentus

Based on our co-pelleting data, we hypothesized that FzlA from *A. tumefaciens* and *R. parkeri* should localize to the site of division in *C. crescentus* since they bind CcFtsZ. We therefore attempted to visualize xylose-induced mNeonGreen (mNG)-FzlA from *C. crescentus*, *A. tumefaciens*, and *R. parkeri* in WT *C. crescentus* to assess localization of FzlA. However, in a WT background we only observed midcell localization of mNG-CcFzlA, while mNG-RpFzlA and mNG-AtFzlA were diffuse. We reasoned that CcFzlA may be outcompeting the FzlAs from *R. parkeri* and *A. tumefaciens* for FtsZ binding, limiting their midcell recruitment.

We therefore sought to localize mNG-FzlA homologs in a *C. crescentus* strain that has *fzlA* deleted. We constructed strains containing xylose-inducible mNG fusions to each of the three *fzlA* homologs in a background in which FtsW and FtsI are hyperactivated, allowing us to delete the native copy of *CcfzlA* (*ftsW**I\** Δ*fzlA*). The *ftsW**I\** Δ*fzlA* strain has a phenotype in which the cells twist around the site of division, resulting in “S” shaped cells prior to division, and exhibit mild elongation, but are otherwise viable and grow similar to WT (8). Each strain was induced with xylose for 1 hour and then imaged (Fig. 6A). We tuned the inducer concentration for each strain to ensure that each mNG fusion was produced to roughly equal levels (Supp. Fig. 3).

**Figure 6.**
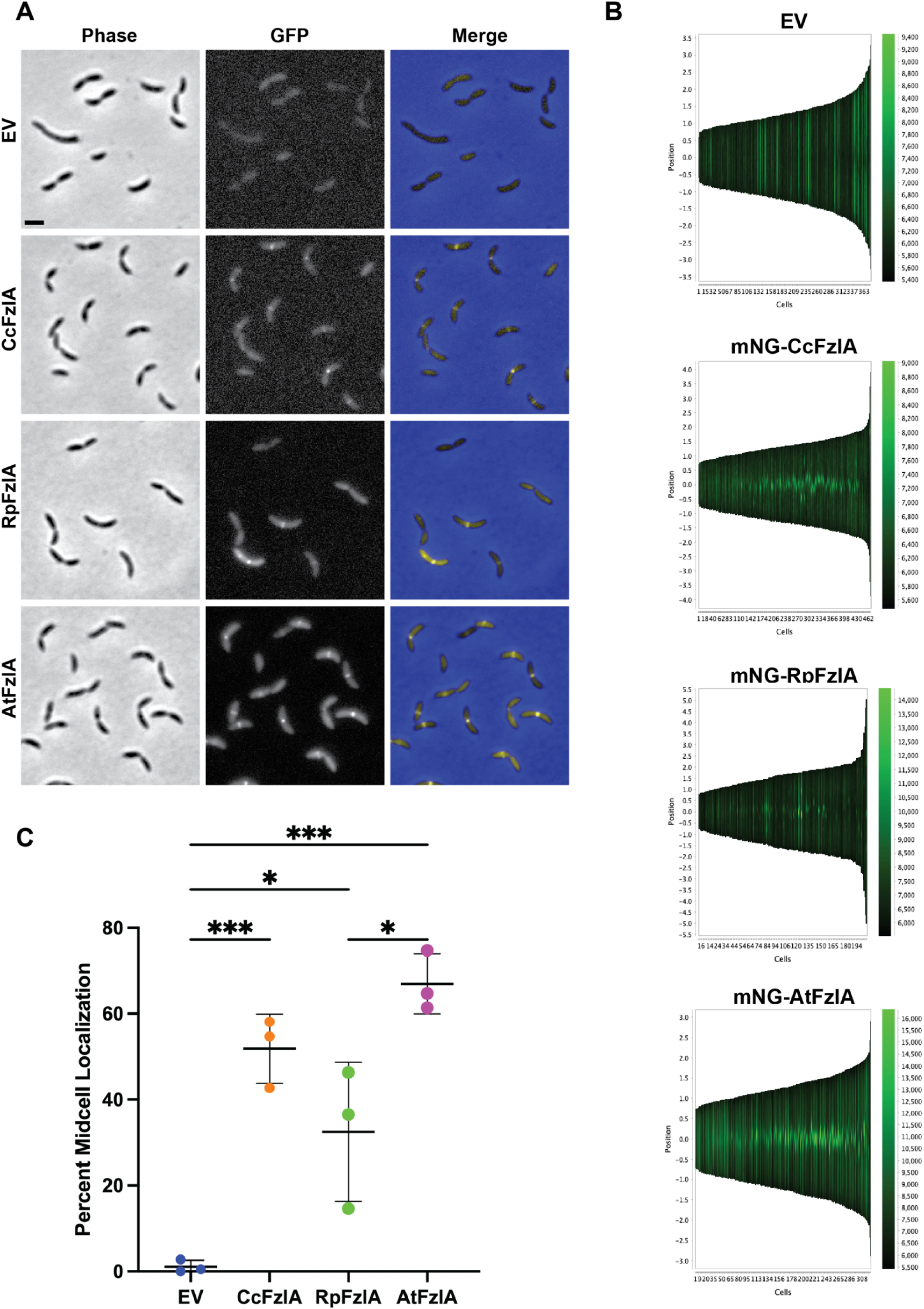
RpFzlA and AtFzlA are able to localize to midcell in *C. crescentus.* **A.** Representative phase-contrast, epifluorescence, and merged images of mNG-FzlA from Cc, Rp, and At expressed in *C. crescentus* under xylose induction. The *mNG-fzlA* fusions were induced with different xylose concentrations for 1 h before imaging to ensure equal protein abundance for the three species. The following concentrations were used: CcFzlA 0.00003% xylose; RpFzlA 0.3% xylose; AtFzlA 0.003% xylose; EV 0.3% xylose. Scale bar = 2 µm. **B.** Representative demographs showing mNG intensity as a function of cell length from CcFzlA, RpFzlA, and AtFzlA mNG fusions. **C.** Graph showing the percentage of cells displaying mNG midcell localization using FzlAs from three alphaproteobacteria. Each condition was performed in triplicate. For each quantification, a one-way ANOVA with Tukey’s multiple comparison test was performed to assess differences between the different conditions tested. *P < 0.0332, ***P < 0.0002.

As expected, we did not observe midcell enrichment for the empty vector control producing cytoplasmic mNG. In contrast, mNG-FzlA from each of the three species localized to midcell in a fraction of cells (Fig. 6A,B). Demograph analysis indicated that each mNG-FzlA was enriched at midcell in stalked and pre-divisional cells, though mNG-RpFzlA appeared to localize to midcell less frequently than the other fusions (Fig. 6B). We determined the fraction of cells with midcell mNG-FzlA and found that, indeed, while mNG-CcFzlA and mNG-AtFzlA localized to the division site in 50% and 65% of cells, respectively, RpFzlA was observed at midcell in only 30% of cells (Fig. 6C). From this analysis we conclude that FzlA from *R. parkeri* and *A. tumefaciens* can properly localize in *C. crescentus* in the absence of CcFzlA, providing further evidence that binding to FtsZ is conserved across species.

### *A. tumefaciens* FzlA partially complements loss of CcFzlA in *C. crescentus*

When we performed our localization experiments in the *ftsW**I\** Δ*fzlA* background we noticed that production of mNG-AtFzlA qualitatively appeared to rescue the twisting and elongation phenotype associated with these cells, suggesting it may complement loss of CcFzlA. We therefore used the strains described in the previous section to quantitatively assess if mNG fusions of *A. tumefaciens* or *R. parkeri* FzlA could rescue the defects associated with loss of CcFzlA in *ftsW**I\** cells. Visual examination of cells suggested that cells producing mNG-AtFzlA resembled those producing mNG-CcFzlA, while cells producing mNG-RpFzlA more closely resembled the strain bearing the EV control (Fig. 6A). We performed principal component analysis of cell shape using Celltool to analyze variations in cell shape across populations (20). For our analysis we focused on the shape modes that primarily reflect cell length (Shape Mode 1) and C-vs S-shape (Shape Mode 3) (Fig. 7A,B). Indeed, consistent with our qualitative observations, cells producing mNG-CcFzlA or mNG-AtFzlA were shorter than those bearing the empty vector, suggesting complementation, whereas those producing mNG-RpFzlA resembled the EV control (Fig. 7A). Similarly, cells producing mNG-CcFzlA or mNG-AtFzlA were primarily C-shaped, whereas S-shaped cells accumulated in cells producing mNG-RpFzlA or bearing the empty vector control (Fig. 7B). The overall distinctions between these strains were most readily visualized when we plotted Shape Mode 3 against Shape Mode 1 (Supp. Fig. 4). Collectively, these data indicate that mNG-AtFzlA is able to rescue the morphological phenotypes associated with loss of *CcfzlA* in the *ftsW**I\** strain, but that mNG-RpFzlA is not (Fig. 7A,B).

**Figure 7.**
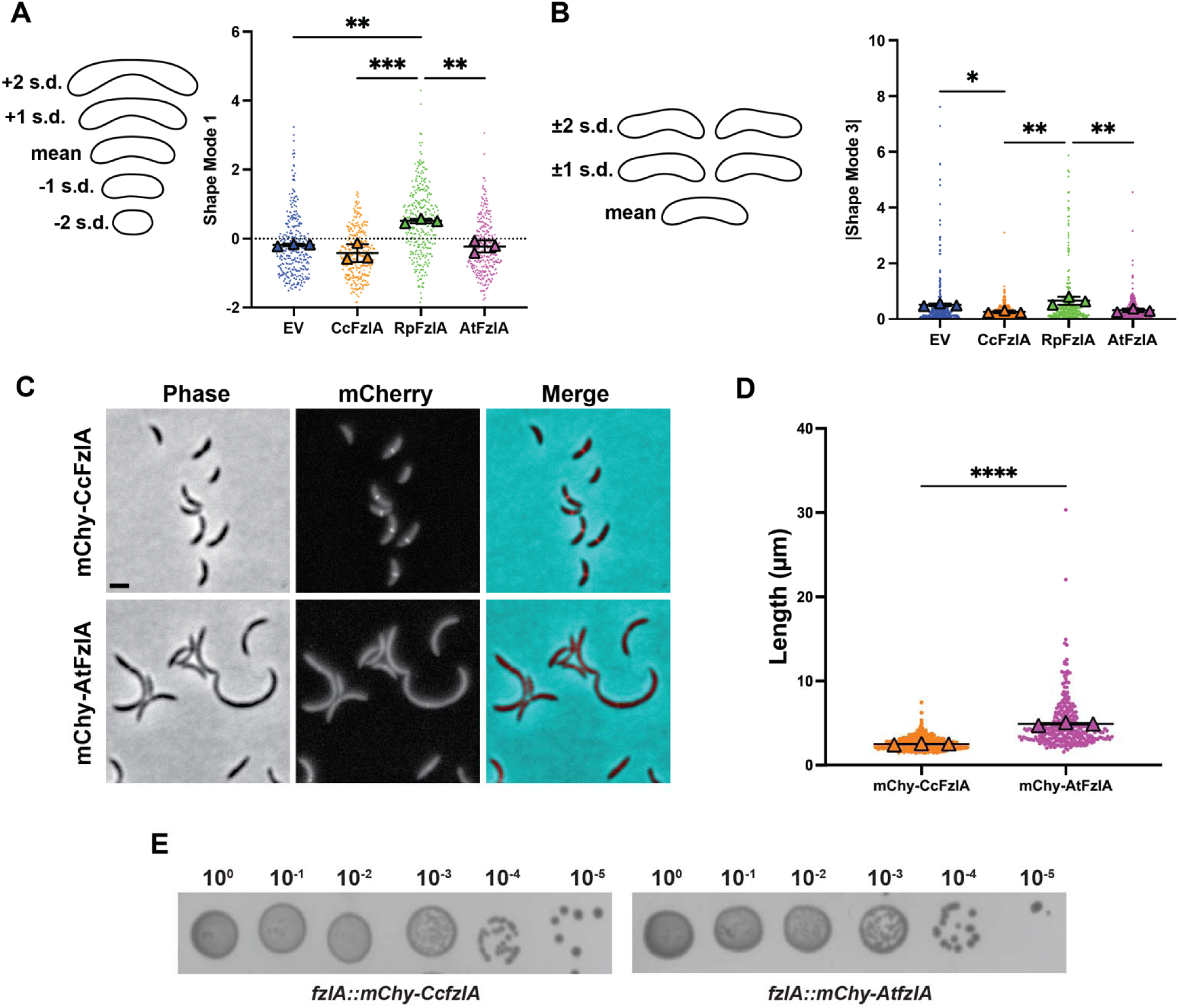
AtFzlA restores cell morphology in *C. crescentus* in the absence of CcFzlA. **A-B.** Principal component analysis (PCA) of cell shape in cells producing mNG-FzlA fusions from different alphaproteobacteria. For each of the shape modes, the mean cell contourL±1 or 2 standard deviations (s.d.) is shown (left). **A.** Shape mode values for shape mode 1 (reflects cell length) are plotted for cells in each strain. A Kruskal-Wallis test with Dunn’s multiple comparison was used to assess differences between the different conditions tested. **P < 0.0021, ***P < 0.0002. **B.** Absolute value of shape mode values for shape mode 3 (reflects C-vs S-shape) are plotted for cells in each strain. A one-way ANOVA with Tukey’s multiple comparison test was performed to assess differences between the different conditions tested. *P < 0.0332, **P < 0.0021. **C-E.** Strains producing either mChy-CcFzlA or mChy-AtFzlA from the *fzlA* locus as the only copy of FzlA in WT *C. crescentus* cells. **C.** Representative phase contrast, epifluorescence, and merged images of *C. crescentus* cells producing either mChy-CcFzlA or mChy-AtFzlA from the *fzlA* locus. Scale bar = 2 µm. **D.** Cell length of each strain described above with the average of each replicate shown (triangles). Cell lengths were quantified using the MicrobeJ program in FIJI. An unpaired t-test was performed to assess differences in the average length between the two strains. ****P < 0.0001. **E.** Representative images of spot dilutions are shown. Each strain was diluted to the same OD and then serially diluted five times on solid PYE.

We next sought to determine whether RpFzlA and/or AtFzlA can localize and function in a WT *C. crescentus* background in place of CcFzlA. We previously made an mCherry-CcFzlA (mChy-FzlA) fusion that was functional when produced from the *fzlA* locus as the only copy of FzlA in the cell. We therefore attempted to introduce genes encoding mChy-AtFzlA or mChy-RpFzlA into *C. crescentus,* replacing *CcfzlA* at the native *fzlA* locus function in a WT background (Fig. 7C). While we were able to obtain strains producing *mChy-AtfzlA*, we were not able to isolate any clones with *mChy-RpfzlA* as the only copy of FzlA produced in the cell. This is consistent with our prior observation that mNG-RpFzlA does not restore normal cell shape to *C. crescentus* cells lacking CcFzlA. We assessed morphology and localization of mChy-AtFzlA-producing cells as compared to cells producing mChy-CcFzlA. mChy-CcFzlA localized to midcell, and cells producing mChy-CcFzlA had an average cell length of 2.5 µm (Fig. 7D). In contrast, we observed rare instances of mChy-AtFzlA midcell accumulation, and cell length was increased to 4.9 µm (Fig. 7D). These mChy-AtFzlA cells were viable as evidenced by our spot dilution assays, suggesting AtFzlA is at least partially functional in *C. crescentus* (Fig. 7E). Collectively, our data indicate that RpFzlA can localize but not complement loss of CcFzlA in *C. crescentus* and that AtFzlA can localize and partially complement loss of CcFzlA.

Finally, we asked if CcFzlA or RpFzlA could rescue the division and viability defects associated with depletion of AtFzlA in *A. tumefaciens* (Supp. Fig. 5). Neither RpFzlA nor CcFzlA were able to rescue loss of AtFzlA, however, as seen by failure to support growth on solid media and cell division defects observed by microscopy (Supp. Fig. 5).

## Discussion

In this study we analyzed the conservation of FzlA function in the alphaproteobacteria *A. tumefaciens* and *R. parkeri*, which exhibit distinct lifestyles and morphologies from *C. crescentus*. Through structural modeling and biochemical assays, we discovered that the FtsZ-FzlA interaction is conserved and binding can occur even across different species, suggesting that conserved residues on FtsZ and FzlA mediate this interaction. Accordingly, we observed that AtFzlA and RpFzlA can both localize to midcell in a *C. crescentus* strain with *fzlA* deleted. In this same strain, we found that AtFzlA can rescue the division defects associated with loss of *C. crescentus* FzlA. Our results suggest that the essential function of FzlA is functionally conserved; however, there are likely variations in its regulation and roles from species to species.

FtsZ is broadly conserved, and our data indicate that its interaction with FzlA is conserved in alphaproteobacteria and essential for regulating constriction. While it is known that FtsZ and FzlA directly interact, many of the details of this interaction are still unknown and we lack a high-resolution structure of the FtsZ-FzlA complex. In this study we used a monomer of FtsZ and FzlA to generate a structural model of their interaction using ColabFold that was consistent with our prior biochemical and genetic observations (Fig. 1). Our assessment of interspecies interaction of FtsZ and FzlA indicates that the interaction interface is at least partially conserved across species. This interaction is required for activation of FtsWI but it is unclear how FzlA binding to FtsZ regulates these enzymes. It could be that FzlA must bind FtsZ simply for midcell recruitment and that, once recruited, it is available to signal through FtsK and other divisome proteins to activate FtsWI. More interestingly, perhaps FzlA alters the polymer structure or biophysical properties of FtsZ to control the activation of FtsWI. Under certain *in vitro* conditions, FzlA can alter the curvature of FtsZ polymers (7), which might facilitate mechanical signaling. However, mutants of FzlA that fail to induce curvature *in vitro* are able to function to some extent in cells, suggesting that it is not an absolute requirement for FzlA function in division (9). Further structural studies, including those capturing FtsZ polymers bound to a FzlA dimer, are required to understand if and how the interaction between FtsZ and FzlA impacts their activities or interactions with other partners.

Our *in vitro* assessment showed conservation of the FtsZ-FzlA interaction in *C. crescentus*, *R. parkeri*, and *A. tumefaciens*. However, in cells, the three FzlAs were clearly not equivalent. While RpFzlA and AtFzlA could localize in *C. crescentus*, presumably because of their ability to bind CcFtsZ, efficient midcell localization required the absence of competing CcFzlA (Figs. 3, 6, 7). Under the more rigorous test of complementation of function, RpFzlA failed to complement and AtFzlA complemented loss of CcFzlA only partially. The reciprocal was not true: CcFzlA could not support cell division when produced in *A. tumefaciens*. These observations are perhaps not entirely surprising when one considers how divergent these organisms, and their corresponding FzlA homologs, are. The primary sequences of the three proteins are only 35-42% identical suggesting significant diversification over time. The inability of CcFzlA to complement loss of AtFzlA function, while AtFzlA could function partially in *C. crescentus*, may suggest additional or auxiliary requirements of FzlA specifically in *A. tumefaciens.* Unlike *C. crescentus*, *A. tumefaciens* grows by polar insertion of new peptidoglycan, followed by a transition to midcell PG synthesis for division. This transition may involve FzlA function in a manner not required in *C. crescentus*. Alternatively, or in addition, *A. tumefaciens* lacks the division site regulator MipZ and the mechanism(s) by which it places the Z-ring at midcell are unknown. FzlA may play an additional role in division site selection in *A. tumefaciens* that is not conserved in *C. crescentus*.

FzlA is conserved throughout the alphaproteobacteria, but not beyond. This observation suggests that FzlA fulfills a function in alphaproteobacteria that is either not required or met by another factor in other bacteria. We recently demonstrated that FzlA interacts with FtsK and proposed that this interaction coordinates chromosome segregation with FtsWI activation (10). In many bacteria, nucleoid occlusion systems exist to ensure the chromosome is not ‘guillotined’ as the cell divides. However, to our knowledge, no nucleoid occlusion factors have been identified in alphaproteobacteria. In *C. crescentus,* constriction begins before complete separation of the replicated terminus regions and the formation of separate nucleoids. This may suggest that the FzlA-FtsK interaction replaces the nucleoid occlusion system to ensure genome integrity during division. In agreement with this, we previously showed that depleting the FtsK DNA translocase domain or overproducing FzlA leads to increased DNA damage, suggesting a role for this interaction in coordinating constriction with chromosome segregation (10). Additionally, the FzlA residues involved in the FtsK-FzlA interaction (residues 223, 227, and 228 in *C. crescentus)* are conserved across alphaproteobacteria, suggesting that this interaction is conserved and could play a central role in regulating the division process. Even *R. parkeri*, which has a streamlined genome and is an outlier in terms of the conservation of proteins in the alphaproteobacteria class, has retained both FzlA and FtsK.

The alphaproteobacteria class encompasses bacteria that have a wide variety of shapes and lifestyles. Their growth environments range from soil, freshwater, and marine environments to growth in or on eukaryotic hosts including humans, animals, arthropods, and plants. Because of these diverse growth environments, alphaproteobacteria have evolved unique morphologies that allow them to thrive in their individual environments. With this morphological diversity, it might be expected that different growth and division mechanisms are required. Perhaps surprisingly then, FzlA is conserved exclusively in this diverse group of bacteria, with homologs present in the vast majority of species (exceptions belong to the obligate intracellular family Ehrlichiaceae, which have non-canonical envelope structure), suggesting it is performing an essential role that has been retained throughout evolution. We propose that FzlA is essential for regulating division by coordinating cell wall synthesis with chromosome segregation in this group. Further study of homologs of FzlA in other alphaproteobacteria will help reveal how FzlA performs this essential function.

## Materials and Methods

### *C. crescentus*, *A. tumefaciens*, and *E. coli* growth medium and conditions

*C. crescentus* NA1000 cells were grown at 30°C in peptone-yeast extract (PYE) medium. *A. tumefaciens* C58 cells were grown at 28°C in *A. tumefaciens* Glucose and Nitrogen (ATGN) medium, and Luria-Bertani (LB) medium when noted. *E. coli* NEB Turbo, Rosetta (DE3)/pLysS, DH5α, S17-1 λ *pir*, and XL10 Gold cells were grown at 37°C in LB medium. Solid media included 1.5% (wt/vol) agar. Antibiotic concentrations used in liquid (solid) media for *C. crescentus* were as follows: kanamycin, 5 (25) µg/mL. For experiments involving inducible gene expression of mNG-tagged FzlAs, inducer concentrations were as follows: CcFzlA 0.00003% xylose(wt/vol); RpFzlA 0.3% xylose (wt/vol); AtFzlA 0.003% xylose; EV 0.3% xylose (wt/vol). Induction with xylose occurred for 1 h before imaging. When indicated for *A. tumefaciens*, kanamycin was used at a concentration of 300 µg/ml, isopropyl-β-D-thiogalactopyranoside (IPTG) was used at a concentration of 1 mM and cumate was used at a concentration of 0.1 mM. Antibiotic concentrations used in liquid (solid) media for *E. coli* were as follows: kanamycin, 30 or 50 (50) µg/ml; ampicillin, 50 (100) µg/mL. Strains used in this study are listed in Supplemental Table 1.

### ColabFold structure prediction

Structures for the FtsZ-FzlA interactions were generated using ColabFold (16). The defaults settings for ColabFold were used except for the following: for the model type alphafold2_multimer_v2 was used and 48 recycles was selected for the number of recycles. ColabFold generated 5 models for each interaction and the top model, as ranked by ColabFold, was selected for analysis in this manuscript.

### FtsZ protein purification

CcFtsZ (pMT219) and AtFtsZ2 (pEG1555) were purified using the protocol described for CcFtsZ in Sundararajan and Goley, 2017 (21). *ftsZ* expression was induced in *E. coli* Rosetta(DE3)/pLysS cells bearing *ftsZ* on a pET21a vector. The culture was grown at 37°C to an OD_600_ of 1.0, at which point protein expression was induced using 0.5 mM IPTG at 37°C for 3 h. Cells were pelleted and resuspended in lysis buffer (50 mM Tris-HCl [pH 8.0], 50 mM KCl, 1 mM EDTA, 10% glycerol, DNase I, 1 mM β-mercaptoethanol, 2 mM phenylmethylsulfonyl fluoride [PMSF], 1 complete mini, EDTA-free protease inhibitor tablet [Roche]) and incubated with 1 mg mL-1 lysozyme for 1 h at 25°C for lysis, followed by sonication. Protein was purified using anion exchange chromatography (HiTrap Q HP, 5 mL; Cytiva) followed by ammonium sulfate precipitation (20 to 30% ammonium sulfate saturation, depending on the protein). The precipitated pellet was resuspended in FtsZ storage buffer (50 mM HEPES-KOH [pH 7.2], 50 mM KCl, 0.1 mM EDTA, 1 mM β-mercaptoethanol, 10% glycerol) and was further purified by size exclusion chromatography (Superdex 200 10/300 GL column; Cytiva). Peak fractions were pooled, aliquoted, and snap frozen in liquid nitrogen for long-term storage at -80°C in FtsZ storage buffer.

RpFtsZ (pEG1936) and AtFtsZ1 (pEG1535) were produced as His_6_-SUMO fusions and cleaved to yield untagged proteins. The His_6_-SUMO-FtsZs were produced from a pTB146 expression vector in *E. coli* Rosetta (DE3)/pLysS cells as described above. Cells were harvested by centrifugation at 6000 x g and resuspended in HK300G (50 mM HEPES-KOH [pH 7.2], 300 mM KCl, 10% glycerol) with 20 mM imidazole, snap frozen in liquid nitrogen and stored at −80°C until purification. To purify, resuspensions were thawed quickly and cells were lysed by incubation with 1 mg/ml lysozyme, 2.5 mM MgCl_2_, and DNase I for 45 min at room temperature followed by sonication. Lysate was cleared and filtered as described above. Protein was isolated by nickel affinity chromatography (HisTrap FF 1 ml, Cytiva) and eluted in HK300G with 300 mM imidazole. Fractions containing His_6_-SUMO fusions were verified by SDS-PAGE, combined with ULP1 Sumo protease at a 1:100 (protease:FtsZ) molar ratio, and cleaved by incubation at 30°C for 3.5 h. Cleaved RpFtsZ or AtFtsZ1 was purified away from His_6_-SUMO by gel filtration (Superdex 200 10/300 GL, Cytiva) in HEK50G. Peak fractions were pooled, snap frozen in liquid nitrogen and stored at −80°C.

### FzlA purification

To purify untagged FzlA for biochemical characterization, His_6_-SUMO-FzlA (pEG994, pEG1885, and pEG1886 for CcFzlA, RpFzlA, and AtFzlA, respectively) or FzlA mutant (pEG1124, pEG1121, and pEG1241 for CcFzlA mutants; pEG1923, pEG1924, and pEG1928 for RpFzlA mutants; pEG1925, pEG1926, and pEG1929 for AtFzlA mutants) was expressed in *E. coli* Rosetta(DE3)/pLysS cells. In initial purification attempts, we found that His_6_-SUMO-RpFzlA and His_6_-SUMO-AtFzlA were not cleaved with Sumo protease and noticed that they were three amino acids shorter than CcFzlA (which was cleaved efficiently) at the N-terminus. We reasoned that the shorter N-terminus may limit accessibility to the protease, so we introduced first three residues of CcFzlA between the SUMO tag and the start codon to extend the N-terminus. This restored Sumo protease cleavage.

Each culture was grown at 37°C to an OD_600_ of 0.4, at which point protein expression was induced using 0.5 mM IPTG at 37°C for 3 h. Cells were pelleted by centrifugation at 6000 x g for 10 min at 4°C, then resuspended in 40 mL of buffer A (50mM HEPES-KOH [pH7.2], 300 mM KCl, 20 mM imidazole, 10% glycerol) per 1 L of culture and snap frozen in liquid nitrogen. The cell suspension was thawed and incubated at room temperature for 1 h, after receiving the following additives: 1 mg/ml lysozyme, 2 units/mL DNAse I (New England Biol-abs) and 2.5 mM MgCl_2_. Cells were sonicated, then centrifuged at 15000 x g for 30 min at 4°C. The supernatant was filtered and loaded onto a HisTrap FF 1 ml nickel column (Cytiva). The column was washed first with 100% buffer A, then 3% buffer B (50 mM HEPES-KOH [pH 7.2], 300 mM KCl, 1 M imidazole, 10% glycerol) to remove non-specifically bound proteins. The His_6_-tagged protein eluted on addition of 30% buffer B. Peak fractions were concentrated, then simultaneously incubated with His_6_-ULP protease to cleave the His_6_-SUMO tag and dialyzed overnight into buffer A. The solution was again loaded onto a HisTrap FF 1 ml nickel column and the flow through was collected, concentrated and dialyzed overnight into storage buffer (50 mM HEPES-KOH [pH 7.2], 300 mM KCl, 10% glycerol), before being snap-frozen in liquid nitrogen and stored at -80°C

### Transmission Electron Microscopy (TEM)

TEM to visualize FtsZ polymers was performed as described in Sundararajan et al., 2015 (18). Thawed protein was diluted to 4 µM in polymerization buffer (50 mM HEPES-KOH [pH 7.2], 50 mM KCl, 0.1 mM EDTA) with 10 mM MgCl_2_. Polymerization was induced with addition of 2 mM GTP and reactions were incubated for 15 min prior to spotting on glow-discharged carbon-coated copper grids (Electron Microscopy Sciences, Hatfield, PA). Grids were blotted and stained twice with 0.75% uranyl formate for 2 min. Grids were then dried and imaged at 100,000 x magnification using a Hitachi 7600 TEM (operated at 80 kV) with an AMT XR80 8-megapixel CCD camera (AMT Imaging).

### Phosphate release assay for FtsZ GTPase activity

Phosphate release by GTP hydrolysis was observed using an assay similar to that described in Sundararajan and Goley, 2017 (21). Thawed protein was diluted to 4 µM in polymerization buffer (50 mM HEPES-KOH [pH 7.2], 50 mM KCl, 0.1 mM EDTA (HEK50) or 300 mM HEPES-KOH [pH 7.2], 50 mM KCl, 0.1 mM EDTA (HEK300)) with either 2.5 mM or 10 mM MgCl_2_. 2 mM GTP was added, and reactions were run for 0 to 30 min in 5 min intervals, stopping each reaction by adding to quench buffer (50/300 mM HEPES-KOH [pH 7.2], 50 mM KCl, 21.3 mM EDTA).

Malachite green reagent (SensoLyte MG Phosphate Assay Kit [AnaSpec]) was added to each reaction and incubated for 20 min before measuring absorbance at 660 nm. Values were compared to a standard curve and plotted to determine GTPase rate. The rate for each protein was measured in triplicate.

### Co-pelleting assays to determine FzlA and FtsZ binding

Co-pelleting assays were performed with 4 μM FtsZ and 8 µM FzlA or FzlA variant in polymerization buffer (50 mM HEPES [pH 7.2], 50 mM KCl, 0.1 mM EDTA) with 10 mM MgCl_2_, 2 mM GTP, and 0.05% Triton X-100. All reactions were incubated at 25°C for 15 min prior to centrifugation at 250,000 x g. Pellet and supernatant from each sample were resolved by SDS-PAGE and stained with Coomassie Brilliant Blue. Gels were imaged with a Gel Doc EZ Gel Imaging System (BioRad) and band intensity was quantified using ImageLab (BioRad). Band intensities were used to calculate the percentage of FzlA or FtsZ present in the pellet.

### Phase contrast and standard epifluorescence microscopy and analysis

For *C. crescentus* microscopy, cells in exponential phase of growth were spotted on pads made of 1% agarose resuspended in water and imaged using a Nikon Eclipse Ti inverted microscope (RRID:SCR_021242) equipped with a Nikon Plan Fluor 100X (NA1.30) oil Ph3 objective and Photometrics CoolSNAP HQ2 cooled CCD camera. For fluorescence, an ET-dsRED filter cube was used for mCherry and ET-GFP for mNeonGreen. Images were prepared for presentation in Photoshop (Adobe) by adjusting fluorescence signal to the same levels across samples in each experiment without oversaturating pixels. Demographs and length analyses were performed using the MicrobeJ plugin for FIJI (22). Each analysis was performed in biological triplicate for each strain, and comparisons were evaluated using one-way ANOVA with Šídák’s multiple comparisons test as indicated in corresponding figure legends. The percentage of cells containing mNG-FzlA at midcell was calculated by manually counting total cells and cells that exhibited midcell fluorescence in FIJI. The percentage of cells with midcell localization was measured in triplicate and graphed using Prism.

For *A. tumefaciens* microscopy, cells in exponential phase were spotted on pads of 1% agarose resuspended in ATGN and imaged using a Nikon Eclipse Ti inverted microscope equipped with a Nikon Plan Fluor 100X (NA1.30) oil Ph3 objective with a QImaging Rolera em-c^2^ 1K EMCCD camera.

### Cell shape analysis

For cell shape analysis, binary masks of phase contrast images of log phase cells were loaded into Celltool (20), allowing for creation of cell contours. Following alignment of cell contours (not allowing for reflection), a model of cell shape was created. The standard deviations away from the mean of each shape mode of interest were plotted as single data points or on a two-dimensional graph plotting two shape modes against each other. GraphPad Prism was used to perform statistical analyses to compare population variances in shape modes across strains.

### Immunoblotting

Equivalent OD units of cell lysate were loaded on an SDS-polyacrylamide gel following cell harvest by centrifugation, resuspension in 1X SDS loading dye, and boiling for 5 – 10 minutes. SDS-PAGE and protein transfer to nitrocellulose membranes were followed using standard procedures. Antibodies were used at the following dilutions: mNeonGreen 1:1,000 (Chromotek); MreB 1:10,000(23); HRP-labeled α-rabbit secondary 1:10,000 (BioRAD Catalog #170-6515); HRP-labeled α-mouse secondary (Cell Signaling Technology Catalog #7076S). Clarity Western Electrochemiluminescent substrate (BioRAD Catalog #170-5060) was used to visualize proteins on an Amersham Imager 600 RGB gel and membrane imager (GE Life Sciences/Cytiva).

### Cell Viability Assessment of *A. tumefaciens* via Spotting Assay

Cells were inoculated into 1 mL of ATGN and incubated for overnight growth of 21 h. Each strain was diluted in ATGN to an OD_600_ = 0.1. Afterwards, 6 h of outgrowth allowed for cultures to reach exponential phase growth (OD_600_ = 0.4-0.6). Each strain was then diluted to OD_600_ = 0.1 before being serially diluted in 10-fold increments for a series of 10^0^ through 10^-7^. After diluting, 3 µL of each dilution was pipetted onto solid LB medium with the appropriate selection and induction. Plates were allowed to dry for 1 h at room temperature prior to being transferred to 28°C for 72 h. Plates were imaged with the BioRad ChemiDoc MP imager under white light.

## Acknowledgements

The authors would like to thank members of the Goley and Brown labs for helpful discussions and comments on this manuscript. We thank Gustavo Santiago-Collazo for strain construction and Regis Hallez for MreB antibodies. This work was supported by grants R35GM136221 (to EDG), T32GM144272 (training grant support of IPP and JMB), and IOS1557806 (to PJBB).

---

**Supplemental Figure 1.**
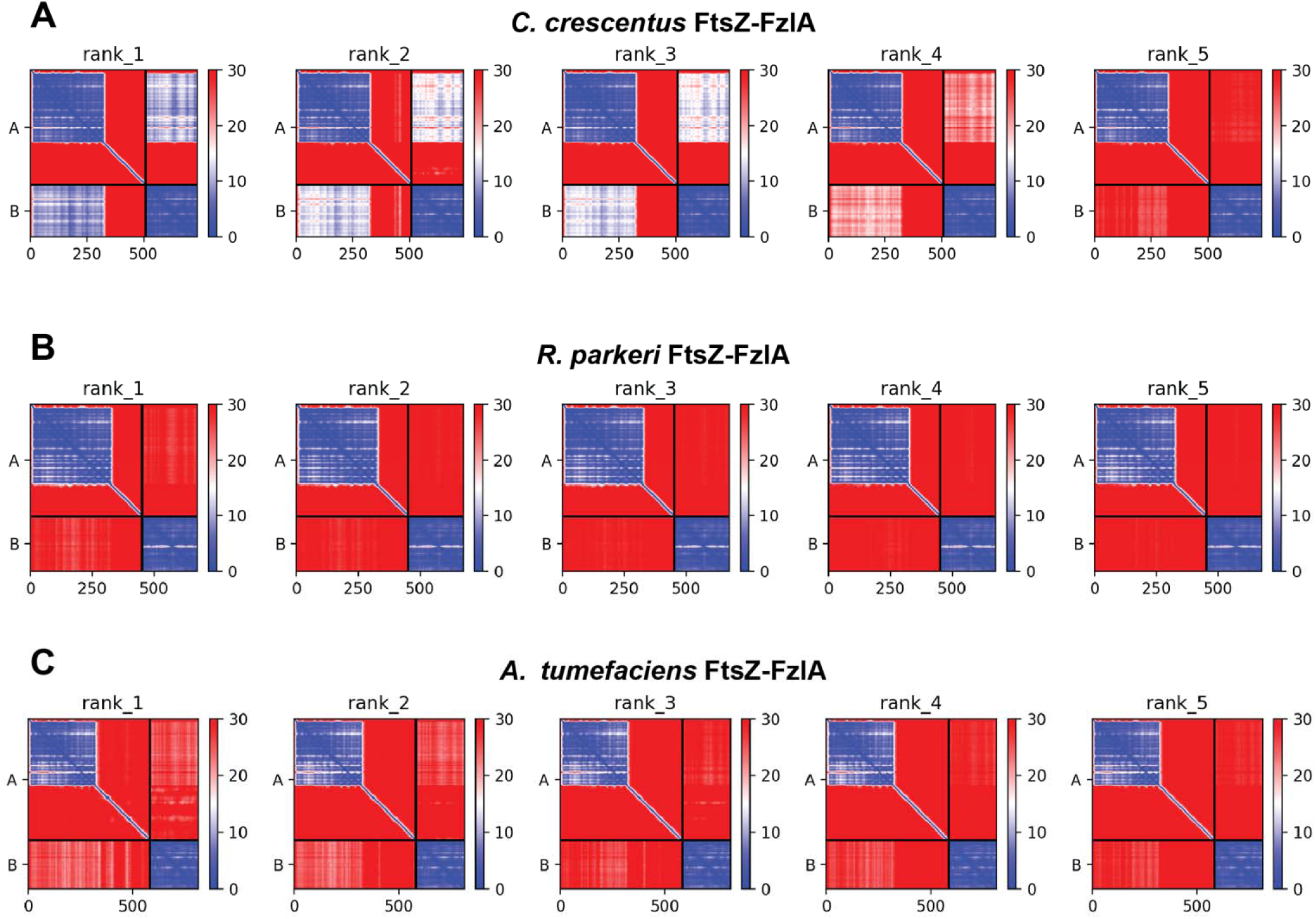
PAE plots of predicted AlphaFold structure for *C. crescentus*, *R. parkeri*, and *A. tumefaciens* FtsZ and FzlA. A-C. PAE plots of the 5 models generated by ColabFold for *C. crescentus* (A.), *R. parkeri* (B.), and *A. tumefaciens* (C.) FtsZ and FzlA. The lower the score, the more confident the prediction. The top-ranked predictions are presented in Figure 1.

**Supplemental Figure 2.**
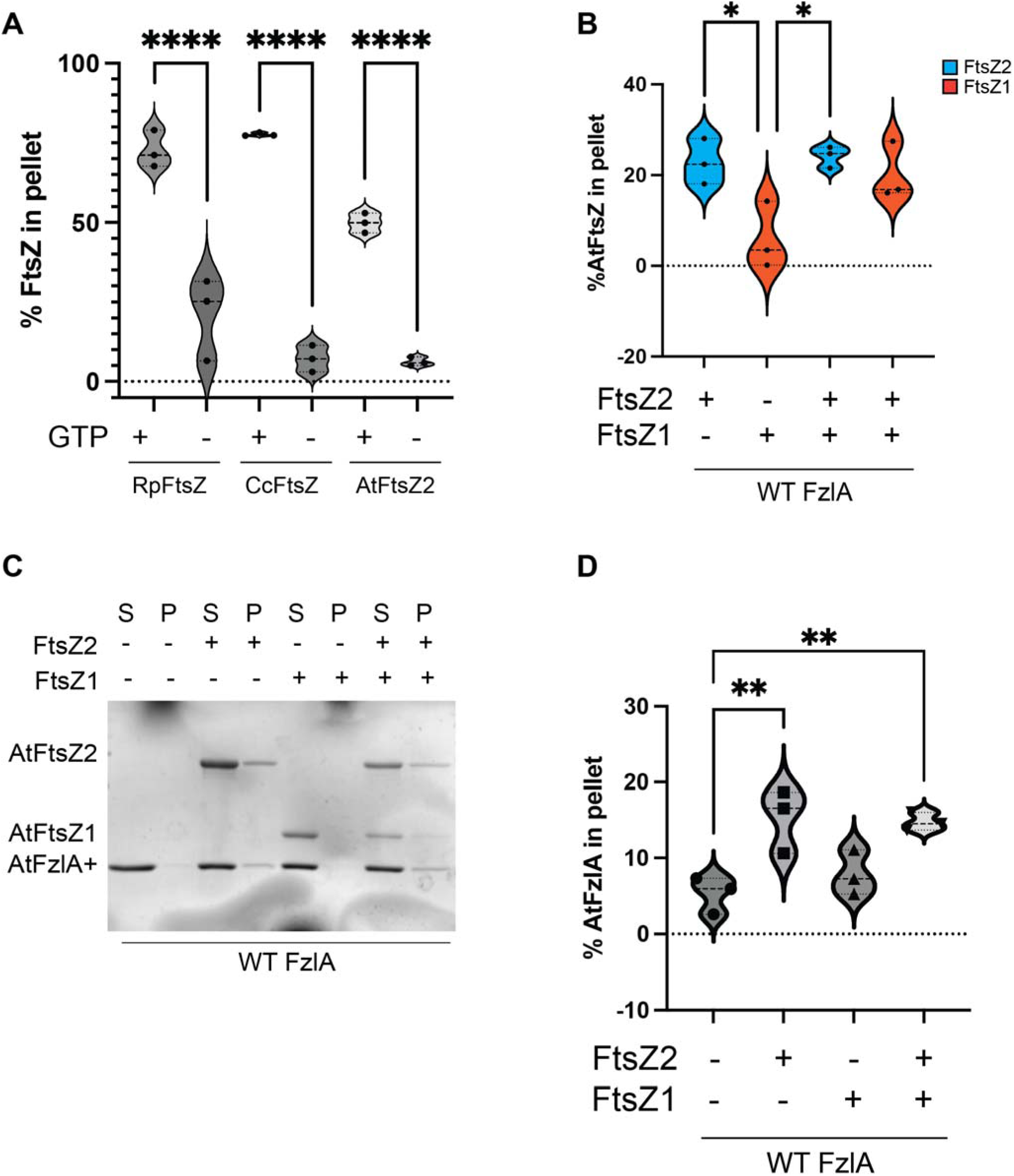
Pelleting properties of alphaproteobacterial FtsZs. **A.** Pelleting of FtsZ from three different alphaproteobacteria with quantification of the amount of FtsZ in the pellet in the presence or absence of GTP. Using the Coomassie blue SDS-PAGE gels in Fig. 3, the amount of FtsZ in the pellet for each condition was quantified and plotted. Each condition was performed in triplicate. **B-D.** Co-pelleting of *A. tumefaciens* FtsZ1 and/or FtsZ2 with AtFzlA**. B.** Quantification of the amount of *A. tumefaciens* FtsZ1 (orange) or FtsZ2 (blue) in the pellet in the presence of AtFzlA. Using the Coomassie blue SDS-PAGE gel in C., the amount of FtsZ in the pellet for each condition was quantified and plotted. **C.** Representative image of the Coomassie blue SDS-PAGE gel from each condition is shown. Using the Coomassie blue SDS-PAGE gels, the amount of *A. tumefaciens* FzlA or FtsZ1/FtsZ2 in the pellet for each condition was quantified and plotted. **D.** Quantification of the amount of AtFzlA in the pellet for each condition shown. All experiments were performed in triplicate. For each quantification, a one-way ANOVA with Šídák’s multiple comparison test was performed to assess differences between the different conditions tested. *P < 0.0332, **P < 0.0021, ****P < 0.0001.

**Supplemental Figure 3.**
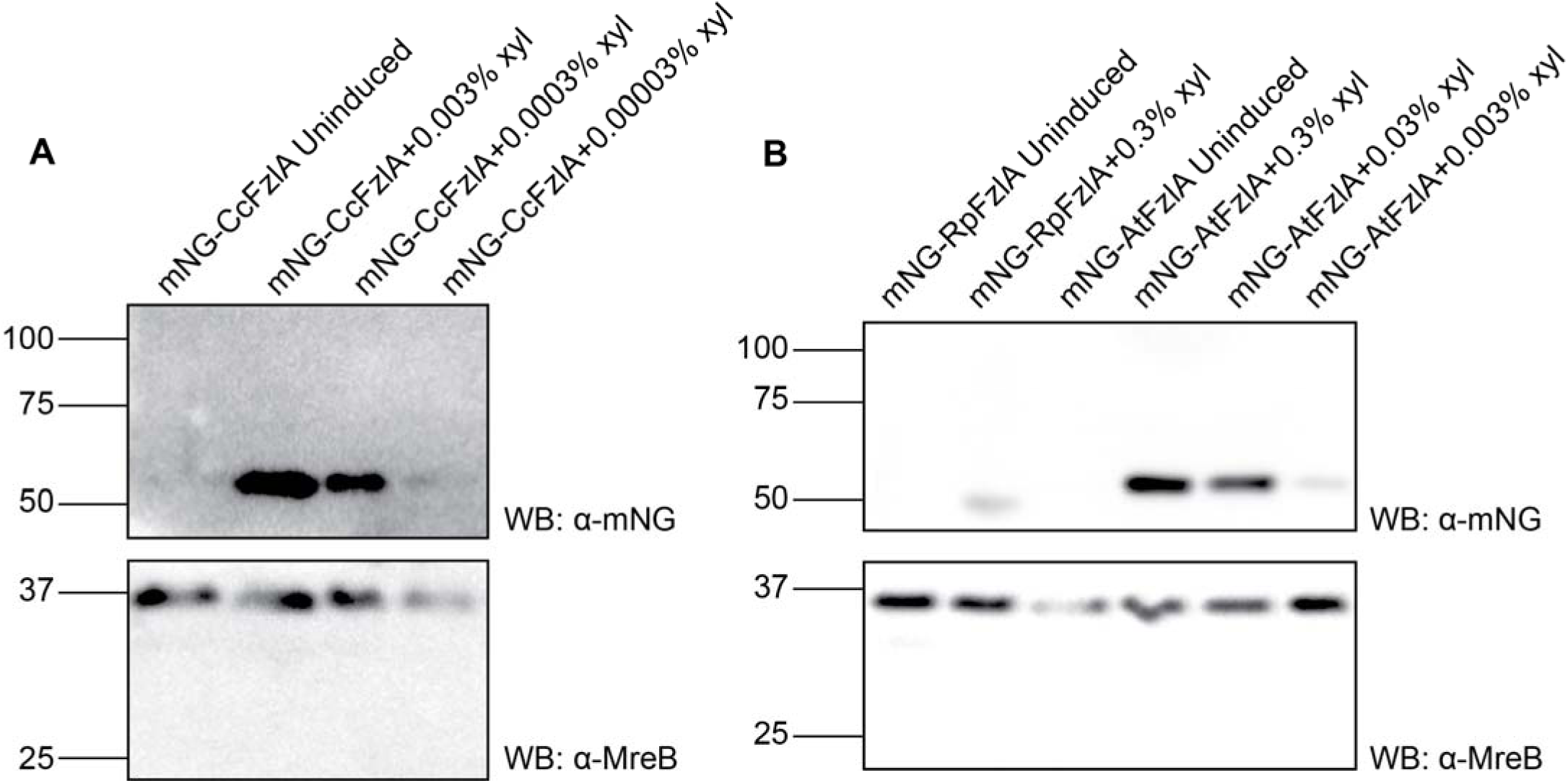
Levels of mNG-RpFzlA, -AtFzlA, and -CcFzlA in *C. crescentus*. **A-B.** Immunoblots using anti-mNG antibody on lysates from *C. crescentus* strains with uninduced or induced mNG-FzlA from different alphaproteobacteria, using varying concentrations of xylose (w/v, xyl). The mNG-FzlA fusions were induced at varying levels for 1 h before samples were collected. MreB was used as a loading control using the anti-MreB antibody.

**Supplemental Figure 4.**
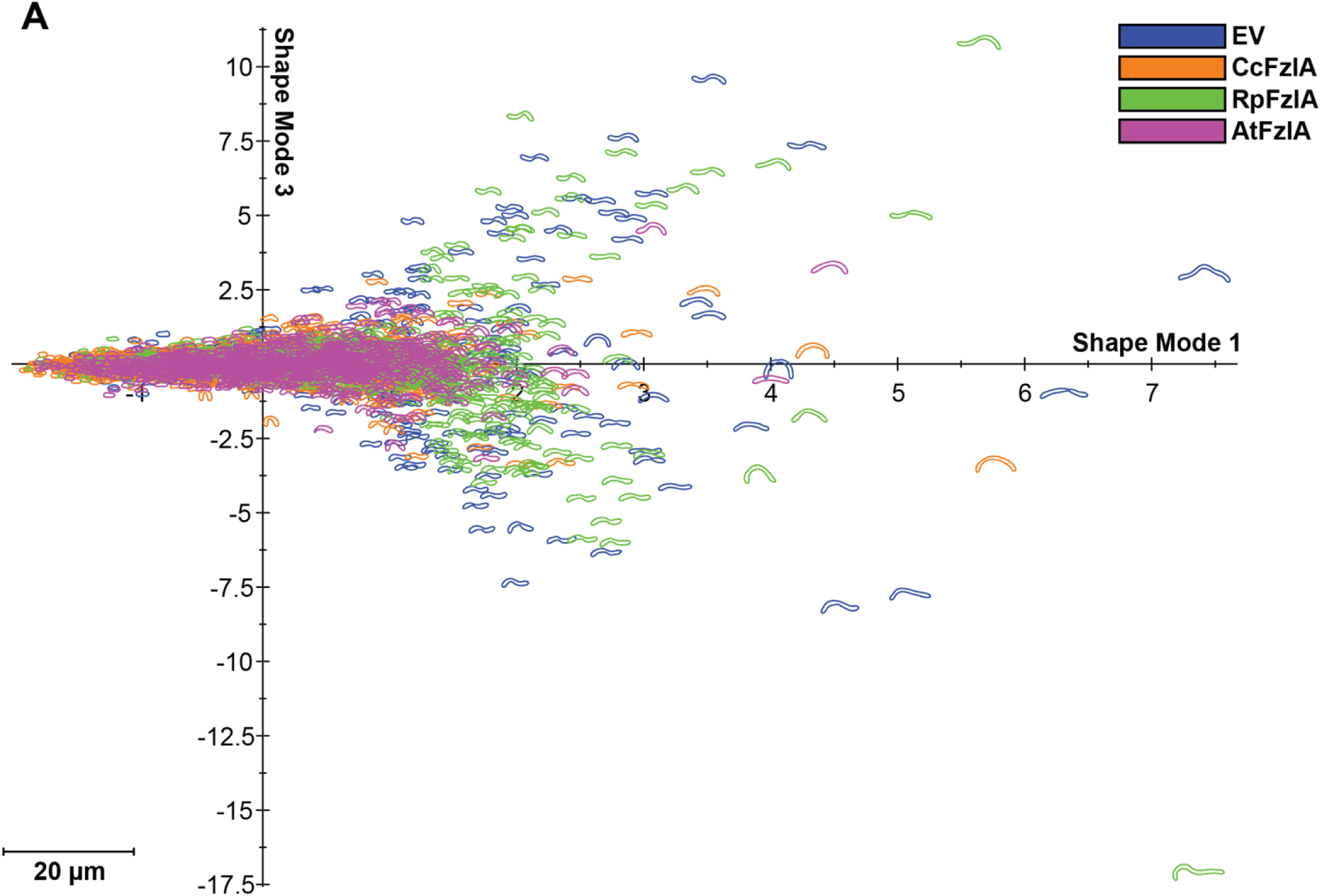
AtFzlA rescues length and twist phenotype in *C. crescentus* lacking FzlA. **A.** Principal component analysis (PCA) of cell shape in cells producing mNG-FzlA fusions from different alphaproteobacteria. Shape mode 1 is plotted against shape mode 3 for each of the strains tested. Description of shape modes can be found in Fig. 7 A,B.

**Supplemental Figure 5.**
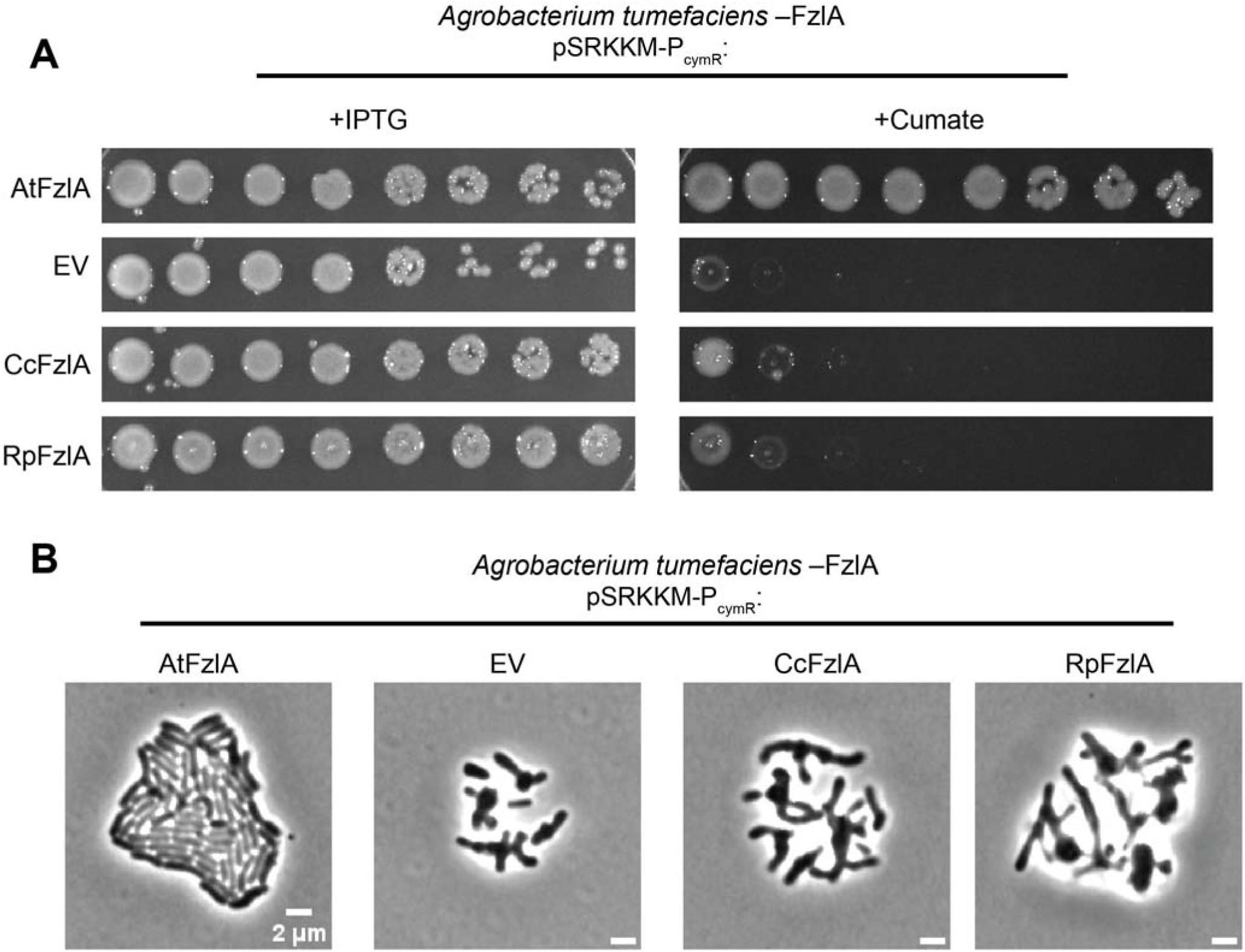
Neither CcFzlA nor RpFzlA can complement the depletion of AtFzlA. **A.** Spotting assays of *A. tumefaciens* in a FzlA depletion background with pSRKKM P*_cymR_*: *AtfzlA*, empty vector, *CcfzlA*, and *RpfzlA*. When grown on LB + IPTG, *fzlA* is expressed from the chromosome. When grown on LB + cumate, the gene indicated to the left is expressed from the plasmid as the only copy of *fzlA*. **B.** Phase contrast images of microcolony formation of AtFzlA depleted cells after 16 h of growth on ATGN + cumate agarose pads to induce the indicated variant of FzlA. EV = empty vector. Scale bar = 2 µm.

## Notes

### Competing Interest Statement

The authors have declared no competing interest.

